# Individual Differences in Neural Decoding Are Stable but Instrument Dependent

**DOI:** 10.64898/2026.09.22.753545

**Authors:** Mohammed Amin Miri, Sara Bonyadian, Rachid Hedjam, Russell Butler

## Abstract

Some participants are consistently easier to decode than others. We asked whether this difference belongs to the person or to the conditions under which the brain is measured. We compared visual decoding in EEG, MEG, intracranial EEG and fMRI, testing classifiers on runs and stimulus identities excluded from training. Within-instrument decoding was reproducible. Attention, behavior and peripheral signals affected recording quality but did not explain most of the stable differences between participants. Anatomy provided one explanation. Source-to-sensor distance strongly predicted modeled signal gain, and its effect was steepest in planar gradiometers. Intracranial recordings provided another: participants with sharper local category selectivity were easier to decode. This association chiefly reflected a smaller response to the nonpreferred category relative to overall visual responsivity. Cross-site prediction was uneven. EEG and MEG expressed the preferred and nonpreferred responses differently. MEG category separation and task-aligned variance described the information available to the decoder. Simultaneous EEG–MEG recordings contained a shared participant component, but external paired cohorts did not yield a common ordering across instruments. Correspondence depended on the measure, the instrument pair and, in one cohort, participant-linked stimulus sets. Repeated observations improved performance in participants with weak single-observation decoding. Stable decoding differences therefore reflect the relation between the individual, the represented distinction and the instrument. They chiefly determine how much information a recording supplies per observation.

## Introduction

Brain recordings differ markedly in the information they yield from one participant to another. Visual information can be decoded from EEG and MEG on a millisecond time scale [1,2,3], from spatial patterns in fMRI [4,5,6], and from intracranial field potentials [7]. Large datasets now relate these measurements to models of visual representation [8,9]. Yet the same procedure may give a clear result in one participant and a weak result in another. These differences affect recording time and complicate comparisons between instruments.

A participant who is repeatedly easy to decode may appear to possess an intrinsically more readable brain. Repetition alone does not establish this. It shows that the ordering is stable under a particular task and method of measurement. Strong cortical selectivity may project poorly to scalp sensors. Favorable anatomy may help one sensor family more than another. A stable difference in attention or physiology may affect several instruments without reflecting a difference in visual representation.

We distinguished three possible sources of variation. The first is behavioral and physiological state. Attention can alter individual trials, and pupil and gaze measurements capture some of these changes [10,11,12]. Eye movements may themselves carry stimulus information [13,14]; motion and physiology may alter BOLD measurements [15,16,17,18]. Their effects on recording quality need not explain why the same participants remain easier to decode. The second source is anatomy: the position and orientation of the cortex relative to the sensors determine how strongly an instrument receives a source [19,20]. The third is the cortical response to the particular distinction being decoded. The instrument converts that response into sensor, voxel or contact patterns, whose separation and variability determine decoding accuracy.

Multimodal recordings allow these explanations to be compared [2,21]. Simultaneous EEG, magnetometers (MAG) and planar gradiometers (GRAD) measure the same neural events through different physical sensitivities. Intracranial EEG (iEEG) measures local cortical responses directly, although electrode placement is determined clinically. fMRI measures a hemodynamic response with different spatial and physiological limits. Paired EEG–MEG–fMRI cohorts then test whether the ordering of the same people survives a change of instrument.

Our working scheme separated effects shared across a person’s measurements from effects specific to the represented distinction, the instrument, or their combination. This was a conceptual division, not a causal model. We asked where predictive information appeared as anatomy and local responses were converted into measured patterns and, finally, decoding accuracy. Runs, stimulus identities and participants were kept separate where required by each test.

The study addressed four questions. Are decoding differences stable, and are they explained by attention or peripheral signals? Does anatomy make the same source more accessible to some instruments than to others? Is local cortical selectivity related to decoding, and how do non-invasive instruments express it? Finally, do simultaneous and externally paired recordings support a common ordering of participants across instruments?

## Results

### Decoding differences are stable and are not chiefly explained by attention

Face–object decoding was highly reproducible within instruments (Fig. 1). Mean balanced accuracy was 0.652 for EEG (n = 95), 0.701 for MAG (n = 100), 0.762 for GRAD (n = 100), 0.671 for iEEG (n = 33), and 0.762 for the exactly reproduced clean fMRI benchmark (n = 102).

**Figure 1.**
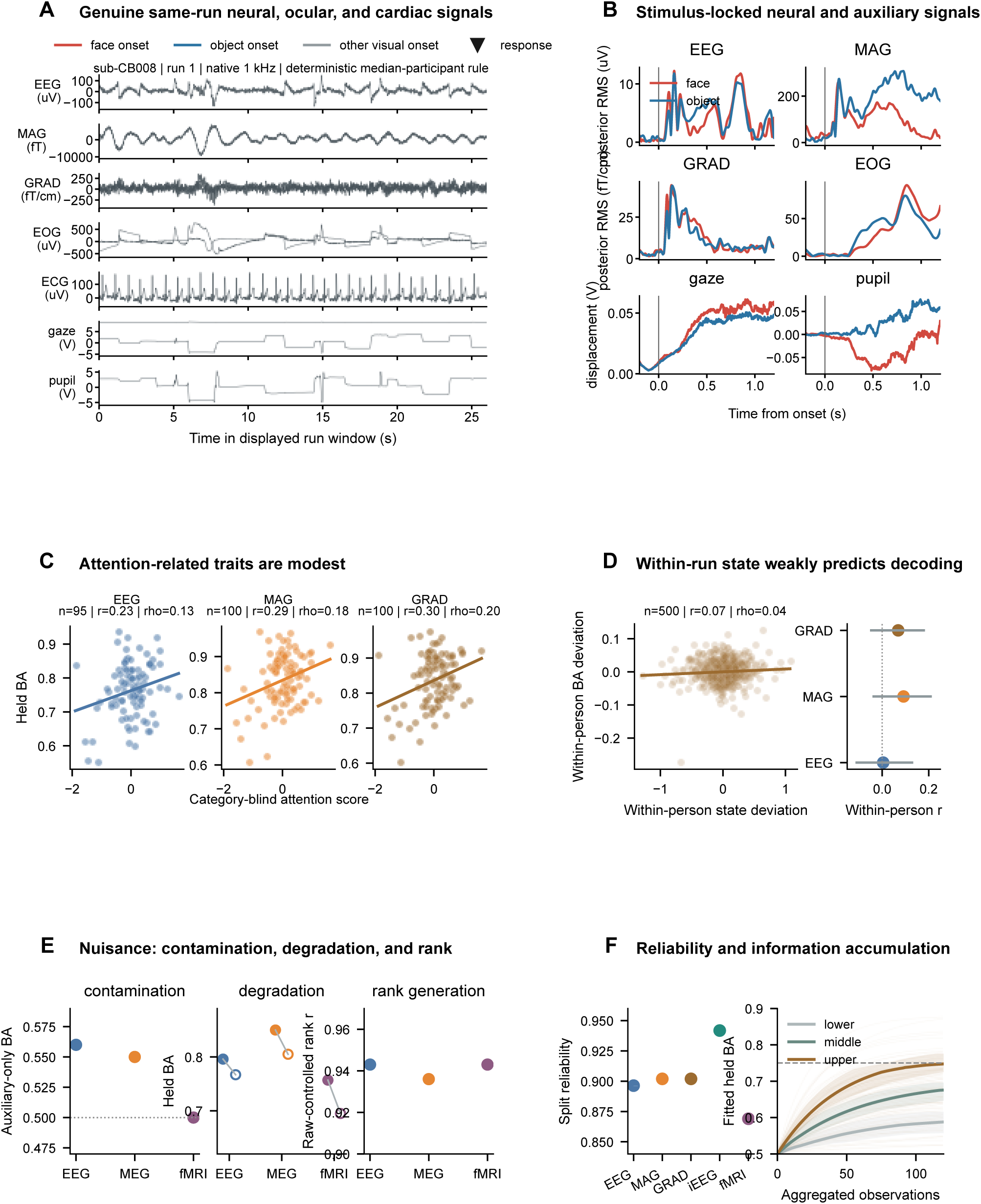
Stable decoding differences and the effects of state. **a. Continuous recordings.** An uninterrupted 26-s segment from run 1 of COGITATE participant sub-CB008. The participant was selected as nearest the joint EEG/MAG/GRAD median, not by visual appearance. Every sample in the original 1-kHz window is retained. Rows show a fixed posterior EEG channel, a posterior MAG channel, a posterior planar-GRAD pair summary, horizontal and vertical EOG, ECG, analog gaze and analog pupil. Vertical markers indicate face, object and other visual onsets; triangles mark recorded responses. Units are native to the recording. Gaze and pupil are uncalibrated analog voltages. **b. Evoked responses.** Face- and object-locked responses from the same participant across five runs, calculated at the original sampling rate. EEG, MAG and GRAD show the RMS of the category-averaged posterior sensor pattern; EOG shows channel RMS. Gaze and pupil retain analog-voltage units. The vertical line marks stimulus onset. Panel a is a continuous record; panel b shows averages derived from event-aligned trials. **c. Attention and participant differences.** Category-blind attention scores plotted against held-out balanced accuracy for EEG (n = 95), MAG (n = 100) and GRAD (n = 100). Scores combine available behavior and auxiliary signals within training folds. Points are participants; lines are descriptive fits. EEG r = 0.23 and ρ = 0.13; MAG r = 0.29 and ρ = 0.18; GRAD r = 0.30 and ρ = 0.20. **d. Within-person state.** Left, run deviations from each participant’s mean state and decoding score across 500 participant-runs (r = 0.07; ρ = 0.04). Right, modality-specific within-person correlations with 95% confidence intervals. **e. Peripheral signals.** Auxiliary-only decoding tests contamination by category information. Paired points show the change in neural balanced accuracy after nuisance control. Raw-to-controlled score correlations test preservation of participant ordering. Dotted lines mark chance where appropriate. Historical fMRI nuisance summaries are retained in the original display as provenance, but were not newly validated by the trial-definition audit. **f. Reliability and accumulation.** Left, archived split-partition reliabilities for EEG, MAG, GRAD, iEEG and fMRI. The fMRI point belongs to the historical measure, not a newly estimated reliability for the clean benchmark. Right, fitted GRAD accumulation curves: thin lines show participants and thick lines summarize lower, middle and upper decoding strata. The dashed line marks balanced accuracy 0.75.

Spearman–Brown split reliabilities for the archived measures ranged from 0.869 to 0.942. The fMRI reliability belongs to the historical measure, rather than to a new reliability estimate for the clean benchmark. Reliable between-participant standard deviation ranged from 0.061 balanced-accuracy units in EEG to 0.127 in iEEG.

The differences depended on what was decoded and where it was measured. Participant performance was strongly related across face–object and four-category decoding, but less strongly across position and duration. Posterior EEG exceeded anterior EEG by 0.0817 balanced-accuracy units and performed better in 92 of 95 participants. Visual and core-posterior iEEG contacts outperformed nonvisual contacts. Visual and fusiform fMRI decoding reached 0.757 and 0.759, respectively, compared with 0.536 in frontal cortex. Adding frontal measurements did not improve the stronger visual signal.

We first inspected continuous neural and auxiliary recordings from the same run, before comparing decoder outputs (Fig. 1a,b). The example contains every sample in a fixed 26-s segment of the original 1,000-Hz recording, with EEG, MAG, GRAD, horizontal and vertical EOG, ECG, analog gaze, analog pupil, stimulus onsets and button responses. The participant was chosen by proximity to the joint median EEG/MAG/GRAD decoding score, not by the appearance of the traces. Face- and object-evoked responses were then calculated at the original sampling rate across all five runs. Trials were averaged before sensor root-mean-square (RMS) amplitude was calculated. These records show the signals entering the analysis; the comparisons of participants use the full cohort.

Attention-related differences were modest (Fig. 1c,d). The plotted category-blind attention scores correlated with decoding at r = 0.23 in EEG (Spearman ρ = 0.13), 0.29 in MAG (ρ = 0.18), and 0.30 in GRAD (ρ = 0.20). Across 500 participant-runs, deviations in state from a participant’s mean were weakly related to deviations in decoding (r = 0.07; ρ = 0.04).

We combined behavior, reaction-time consistency, gaze stability, pupil validity and response, blinks, EOG, cardiac measures and artifacts in models fitted within training folds. Adding these state measures to acquisition variables did not improve held-participant prediction in EEG, MAG or GRAD: the increments were negative. The iEEG model produced positively correlated predictions but was severely miscalibrated. Lagged and prestimulus analyses likewise showed that favorable state improved trial quality without selectively rescuing weak participants or removing their stable ordering. The available run-level data did not support a single trait–state variance decomposition for every modality.

Peripheral signals affected absolute accuracy more than participant ordering (Fig. 1e). EOG carried modest category information. Linear and nonlinear nuisance controls reduced mean EEG decoding from 0.796 to 0.767 and 0.781, and MEG decoding from 0.850 to 0.805 and 0.825. Nevertheless, raw and controlled participant scores remained closely related: r = 0.943 and 0.965 for EEG, and 0.936 and 0.960 for MEG. Ocular signals contained category information, and nuisance control reduced neural decoding, but participant ordering remained largely unchanged. Historical fMRI nuisance and state analyses used a different measure whose reproduction remains unresolved; their status is detailed in Methods.

Additional observations improved accuracy for most participants with weak single-observation performance (Fig. 1f). In GRAD, category separation predicted the rate of improvement in both directions of cross-site testing (r = 0.380 and 0.502). Recurrence, variance along the discriminating axis, nuisance measures and modeled gain added no transferable prediction beyond separation.

Fitted counts of observations needed to reach an absolute accuracy criterion were unstable when the estimated asymptote fell below that criterion. The relative result was clear: participants differed in how quickly evidence accumulated. Weak decoding usually meant less information per observation, rather than no information.

### Anatomy changes what each instrument can measure

The same cortical source produced different sensor fields in different participants. We passed an identical unit-energy bilateral visual source through each participant’s anatomy and forward model (Fig. 2a). The examples were selected at specified percentiles of modeled gain, independently of decoding accuracy. Differences in their T1-derived pial geometry changed the GRAD field even though source energy was fixed.

**Figure 2.**
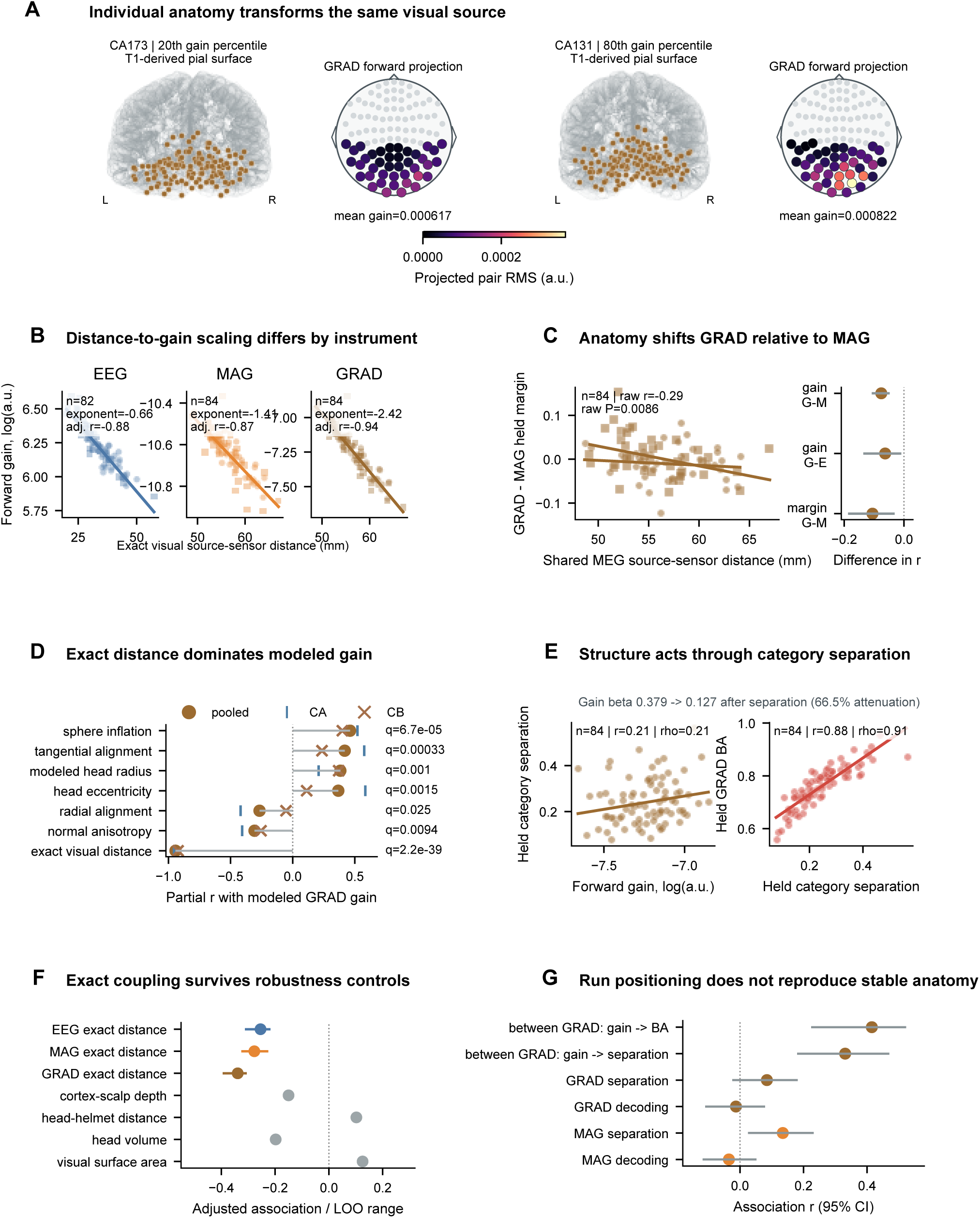
Anatomy changes the signal received by the instrument. **a. Identical sources, different projections.** T1-derived pial anatomy and GRAD forward projections for participants nearest the 20th (CA173) and 80th (CA131) percentiles of visual forward gain. Displayed mean gains are 0.000617 and 0.000822, respectively. Selection used modeled gain, not decoding. An identical unit-energy bilateral visual source was passed through each participant’s forward operator. GRAD pairs are shown as root-sum-square magnitudes at unique locations on a shared nonnegative scale. These are modeled fields, not measured activation. **b. Distance and gain.** Exact bilateral visual source-to-sensor distance plotted against gain for EEG (n = 82), MAG (n = 84) and GRAD (n = 84). Shapes identify sites and lines show pooled descriptive fits. The displayed rounded log–log exponents are −0.66, −1.41 and −2.42; adjusted correlations are −0.88, −0.87 and −0.94. Full estimates are given in Results. **c. Relative GRAD sensitivity.** Left, shared MEG source-to-sensor distance against the within-participant GRAD-minus-MAG held margin (n = 84; raw r = −0.29, P = 0.0086). Right, paired differences in distance associations for gain and held margin, with bootstrap 95% confidence intervals. **d. Structural associations with gain.** Pooled and site-specific adjusted associations between seven anatomical measures and modeled GRAD gain. Exact distance dominates. q values use the prespecified family correction. The displayed q annotations are 2.2 × 10^−39^ for exact distance, 0.0094 for normal anisotropy, 0.025 for radial alignment, 0.0015 for head eccentricity, 0.001 for modeled head radius, 0.00033 for tangential alignment and 6.7 × 10^−5^ for sphere inflation; Results retain the reported narrative estimates. **e. Gain, separation and decoding.** Left, gain against cross-fitted GRAD category separation (n = 84; displayed r = 0.21, ρ = 0.21). Right, separation against held-out balanced accuracy (n = 84; displayed r = 0.88, ρ = 0.91). Adding separation reduced the locked standardized gain–decoding coefficient from 0.379 to 0.127, or 66.5%. This is an attenuation analysis, not causal mediation. **f. Anatomical controls.** Exact-distance associations are compared with cortex-to-scalp depth, head-to-helmet distance, head volume and visual surface area. Colored estimates show exact coupling; muted estimates show the controls. **g. Between-person and within-session effects.** Ordinary changes in position between runs do not reproduce the stable between-participant association. Forward quantities describe measurement physics, not the strength of cortical representations.

Source-to-sensor distance predicted modeled gain in all three sensor families, most steeply in planar gradiometers (Fig. 2b). The log–log exponents were −0.659 for EEG, −1.411 for MAG and −2.421 for GRAD. Held-out linear distance models explained R² = 0.838, 0.713 and 0.861, respectively. Distance also predicted decoding after adjustment for age, sex and acquisition: partial r = −0.254 for EEG (permutation P = 0.0104), −0.277 for MAG (P = 0.0136), and −0.339 for GRAD (P = 0.0023). Modeled gain was weakly related to EEG decoding (r = 0.157), but more strongly related to MAG (r = 0.326, P = 0.0035) and GRAD (r = 0.385, P = 0.0002). Source alignment was significant in MAG and GRAD, but not EEG (Supplementary Results).

Seven anatomical measures were associated with GRAD gain after correction for multiple testing (Fig. 2d). Exact source-to-sensor distance dominated (partial r = −0.943, q = 2.25 × 10^−39^). The remaining associations were tangential alignment (r = 0.419, q = 3.29 × 10^−4^), radial alignment (r = −0.267, q = 0.025), visual-normal anisotropy (r = −0.307, q = 0.009), modeled head radius (r = 0.383, q = 0.001), sphere inflation ratio (r = 0.462, q = 6.74 × 10^−5^), and head-shape eccentricity (r = 0.367, q = 0.002). These measures were correlated. Distance dominated at both sites; the secondary shape and orientation effects were less consistent across sites.

Broad anatomical measures were less useful (Fig. 2f and Extended Data). Anatomy-family models gave held R² = 0.036 in EEG and 0.287 in MAG, while the strict common-sample GRAD increment was 0.004. Historical fMRI anatomy models remain in the audit supplement. Lateral-ventricle fraction was a secondary GRAD marker, with adjusted standardized effects of 0.255 for decoding, 0.295 for reproducibility and 0.294 for category separation. Only separation survived correction across the full family (q = 0.042; decoding q = 0.095; reproducibility q = 0.099). The direct association with decoding disappeared after adjustment for separation.

A within-person comparison made the instrument dependence explicit (Fig. 2c). Greater source-to-sensor distance predicted a smaller GRAD-minus-MAG held-margin difference (n = 84; raw r = −0.29; adjusted r = −0.330, P = 0.00234). The difference in adjusted distance correlations was −0.106 (95% CI, −0.187 to −0.031; P = 0.0026), showing greater GRAD sensitivity on the margin measure. The corresponding balanced-accuracy interaction was inconclusive (P = 0.119).

Exact distance remained related to EEG and GRAD decoding after adjustment for orientation. Coarse cortical depth, head-to-helmet distance, head volume and visual surface area gave weak or inconsistent results. Ordinary changes in head position between runs did not reproduce the stable differences between people (Fig. 2f,g). The relevant anatomy was the relation of the source to the instrument, not a general advantage of head size or cortex.

The effect was expressed in the measured category response (Fig. 2e). In the locked GRAD analysis, gain predicted category separation. Adding separation reduced the standardized gain– decoding coefficient from 0.379 to 0.127, an attenuation of 66.5%. Independent reconstruction recovered the same direction and 61.6% attenuation with a different available-participant and control set. These results connect source–sensor coupling, modeled gain and the separation of category responses at the sensors. They describe attenuation of an association, not causal mediation or a change in the underlying cortical representation.

### Local cortical selectivity predicts decoding

Direct recordings showed a local cortical correlate of the decoding differences. The core iEEG analysis included 569 anatomically visual contacts from 29 participants (Fig. 3a,b). We defined category preference and tuning in training runs and identities, then measured the responses on separate observations. Some local populations responded very differently to faces and objects; others responded similarly to both.

**Figure 3.**
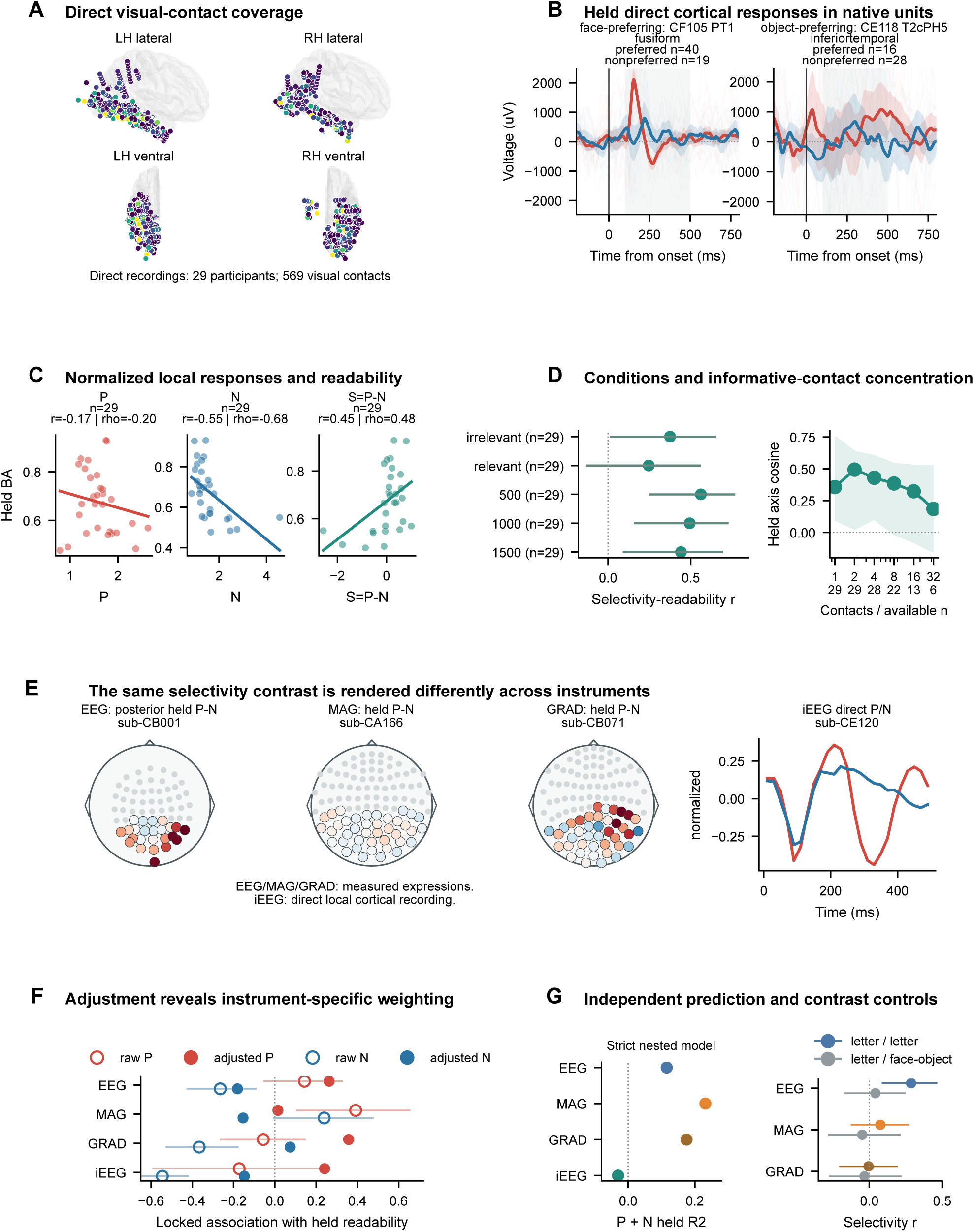
Local selectivity and its expression across instruments. **a. Intracranial coverage.** The 569 anatomically visual contacts from 29 participants, shown on fixed lateral and ventral cortical views. Color indicates cross-fitted tuning. **b. Local responses.** Held-out voltage responses from face-preferring fusiform contact CF105 PT1 (preferred n = 40; nonpreferred n = 19) and object-preferring inferior-temporal contact CE118 T2cPH5 (preferred n = 16; nonpreferred n = 28). Preference and contact selection were defined without the displayed observations. Thin lines show trials, thick lines show means and bands show 95% trial intervals. The vertical line marks onset; voltage is shown in native units. **c. Response components.** Preferred response P, nonpreferred response N and selectivity S = P − N plotted against locked held-out iEEG balanced accuracy (n = 29). P and N are normalized by the held-out all-visual response. Displayed r/ρ values are −0.17/−0.20 for P, −0.55/−0.68 for N, and 0.45/0.48 for S. Lower N denotes a lower nonpreferred response relative to overall responsivity, not necessarily a lower absolute voltage or synaptic inhibition. Training-normalization and absolute-response tests are reported in Methods and Supplementary Methods. **d. Duration, relevance and contact accumulation.** Left, independently reconstructed condition-matched selectivity– decoding correlations, with one observation per participant (n = 29 per condition) and Fisher-z 95% intervals. Right, median held category-axis cosine as training-ranked contacts are added. Bands are participant interquartile ranges, not confidence intervals. Ticks give contact count and available participant count; participants are not extrapolated beyond their available contacts. **e. Measured selectivity across instruments.** The closest valid face–object selectivity displays: posterior EEG for sub-CB001, MAG for sub-CA166, pair-combined GRAD for sub-CB071, and direct iEEG P/N responses for sub-CE120. EEG, MAG and GRAD are sensor-level expressions; iEEG is a direct local recording. Spatial maps use the appropriate measurement stage. The COGITATE fMRI component display was removed. **f. Raw and adjusted associations.** Locked P and N associations with decoding. Open and filled markers distinguish raw and adjusted values. Lines are bootstrap confidence intervals for raw estimates only; adjusted points are descriptive and have no displayed intervals. **g. Prediction and independent contrasts.** Left, strictly nested P + N held-participant R² for EEG, MAG, GRAD and iEEG, with scaling refitted in every inner fold and negative prediction scores retained. These are absolute predictions from the independent model, not increments from the locked adjusted model in panel f. Right, held-run/identity letter/font selectivity correlations with letter/font and separate face/object decoding in EEG (n = 95), MAG (n = 100) and GRAD (n = 100). Same-target intervals are Fisher-z 95% intervals; cross-target intervals use participant bootstraps. No fMRI component estimate is included.

Participants with sharper local tuning were easier to decode (Fig. 3c). Mean top-contact tuning correlated with the locked iEEG decoding measure at r = 0.736 (ρ = 0.759). Independent reconstruction gave an anatomy-adjusted association of r = 0.452 (ρ = 0.364); a complementary participant random-intercept analysis gave r = 0.496 (ρ = 0.413). A model trained at site CE predicted site CF well (held R² = 0.522, r = 0.818), whereas the reverse direction was weak (R² = 0.046, r = 0.307). Stronger archived robust top-contact models used different specifications and are reported separately in the supplement.

The distinction lay chiefly in the response to the nonpreferred category. We divided each selected contact’s response into preferred expression, P, and nonpreferred expression, N, normalized by overall visual responsivity. Preferred expression was unrelated to decoding (r = −0.173; ρ = −0.199). Lower nonpreferred expression was associated with better decoding (r = −0.545; bootstrap 95% CI, −0.753 to −0.419; ρ = −0.682; permutation P = 0.0005). Selectivity, P − N, was positively associated with decoding (r = 0.451; 95% CI, 0.211–0.663).

The nonpreferred association survived exclusion of contacts above a tenfold amplitude threshold (r = −0.543) and deletion of any one participant (r = −0.644 to −0.506). Participant sub-CG102 was nevertheless influential (Cook’s D = 3.98). The nested component models predicted individual scores poorly: held R² was −0.030 for P + N after strict inner-fold scaling, 0.042 for N alone, and −0.143 for P alone. A strong association therefore did not amount to a well-calibrated prediction model in this small sample.

Normalization mattered. Absolute nonpreferred amplitude was not marginally associated with decoding (n = 29, r = −0.137, P = 0.479). Using a denominator estimated only from training observations preserved the negative association (r = −0.579; ρ = −0.685; bootstrap 95% CI, −0.739 to −0.463). In a joint model of absolute preferred and nonpreferred responses, training-derived overall magnitude and site, the standardized nonpreferred coefficient was −0.900 (HC3 95% CI, −1.288 to −0.511), while the preferred coefficient was 0.008. The result is lower nonpreferred response relative to overall responsivity, not a general reduction in absolute amplitude. These sensitivity tests used the same cohort. They concern measured responses, not synaptic inhibition.

The association was already present at the shortest presentation duration (Fig. 3d). Independently reconstructed selectivity–decoding correlations were 0.561, 0.494 and 0.441 at 500, 1,000 and 1,500 ms, respectively (n = 29 in each condition). It was also present when the stimuli were task irrelevant (r = 0.375, P = 0.045). The relevant-trial estimate was weaker (r = 0.246, P = 0.198). Prolonged viewing and explicit task relevance were therefore not prerequisites, although the two relevance conditions did not each establish significance.

Training-ranked contact accumulation gave median held-axis cosines of 0.357, 0.494 and 0.430 with one, two and four contacts. The available sample was 29 participants at one and two contacts, then 28, 22, 13 and 6 at four, eight, sixteen and thirty-two contacts. This analysis measured how category-axis information was concentrated among contacts; it did not show a monotonic decoding benefit from adding electrodes.

The non-invasive measurements did not give a single preferred/nonpreferred pattern (Fig. 3e–g). The locked EEG analysis showed a weak analogue of the lower-nonpreferred iEEG effect, while MAG showed broader positive expression. In GRAD, the locked adjusted analysis favored preferred expression, but the independent P + N model assigned a larger unique contribution to N. Its N association remained negative after adjustment for separation (r = −0.320). Independently nested P + N models gave held R² = 0.116 for EEG, 0.232 for MAG and 0.175 for GRAD. Thus the GRAD attribution depended on the model. The COGITATE fMRI component analysis was excluded because its all-trial decoding measure did not reproduce; the clean common-decoder benchmark did reproduce exactly (Methods).

Nor did selectivity for one distinction reliably predict decoding of another. With separate runs and identities, letter/font selectivity predicted letter/font decoding in EEG (r = 0.286, P = 0.0049), but not MAG (r = 0.076, P = 0.455) or GRAD (r = −0.005, P = 0.959). Its associations with face/object decoding were small: r = 0.042, −0.049 and −0.033, respectively. The corresponding iEEG cross-contrast association was also unsupported. These controls did not establish a category-general selectivity trait; the nonsignificant cross-contrast tests alone do not establish a statistical difference from the within-contrast effects. Earlier fMRI component controls are archived as withdrawn.

The measured separation of category responses provided the closest link to MEG decoding. Combined category-blind predictors added R² = 0.063 beyond acquisition in EEG and MAG, and 0.303 in GRAD; iEEG prediction was miscalibrated (incremental R² = −2.179). General response accessibility was the strongest upstream family in GRAD, whereas MAG was strongly structured by acquisition. Historical COGITATE fMRI upstream results remain pending reconciliation.

Whole-head RMS amplitude was unrelated to site-adjusted MEG rank (MAG r = 0.052; GRAD r = 0.053). Independently reconstructed category separation, by contrast, correlated with decoding at r = 0.881 in MAG and 0.875 in GRAD. Variance parallel to the discriminating axis added held-out prediction, and global amplitude normalization preserved participant rank. Temporal response profiles recurred strongly within the target contrast, but noise-covariance recurrence and recurrence across independent contrasts did not establish a general readable-brain trait. Separation and decoding summarize related response distributions. They locate the information in the measured response, rather than provide independent evidence for an upstream biological cause.

### Shared participant effects do not give a universal ordering across instruments

Simultaneous EEG–MEG recordings contained a substantial common participant component (Fig. 4d). The locked multilevel decomposition assigned 42.9% of total variance to participant-general effects, 11.1% to participant-by-modality effects, 21.7% to participant-by-task effects and 24.3% to unresolved residual variance. Considering only the general and modality-specific components gave 79.4% general and 20.6% modality specific. These last two percentages exclude task-related and residual variance. A separate common factor accounted for 86.7% of EEG–MEG covariance.

**Figure 4.**
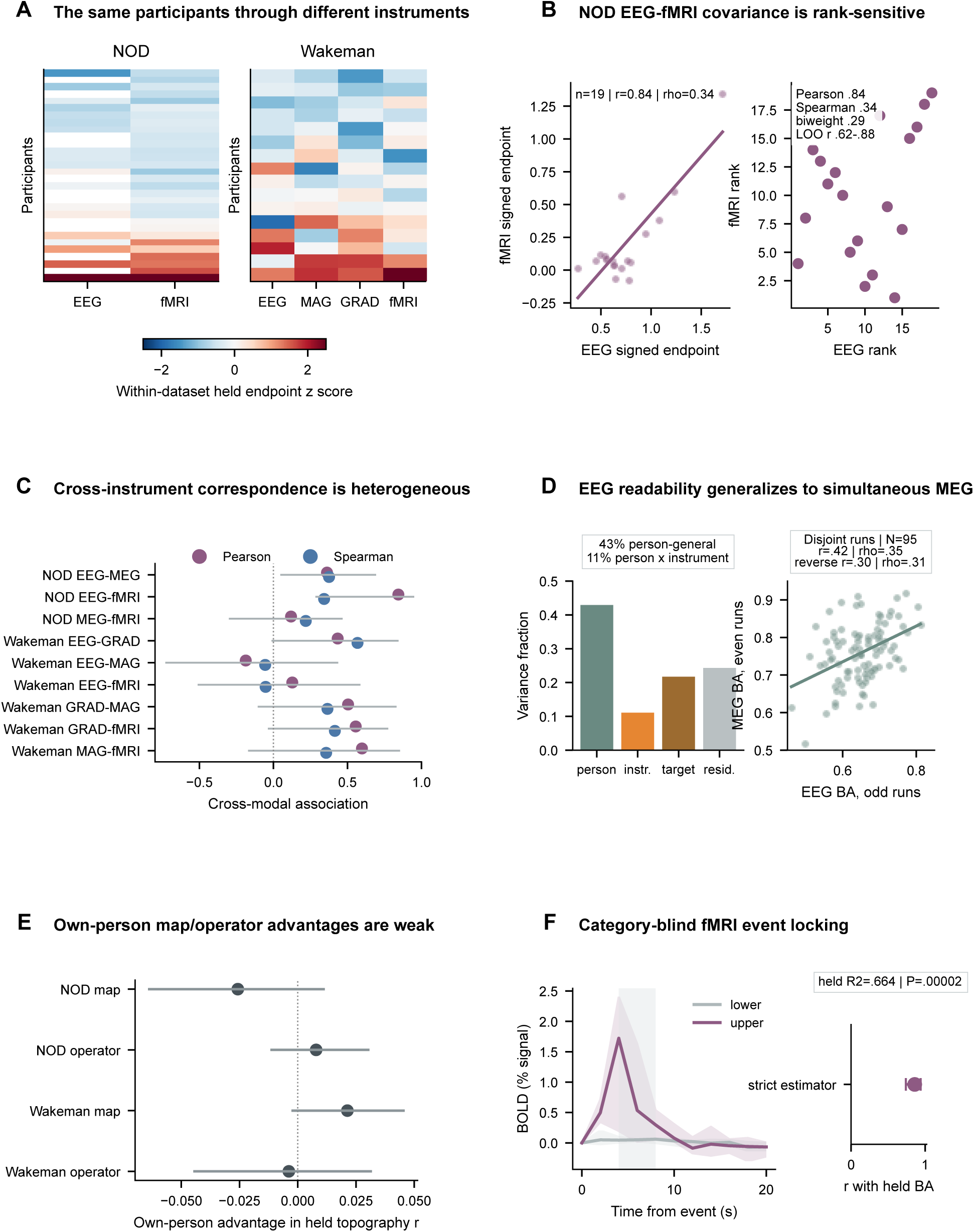
Shared and instrument-specific participant differences. **a. The same participants across instruments.** Held-out score heat maps for NOD and corrected Wakeman–Henson data. Scores are standardized within dataset and modality. Participants are ordered by their mean available standardized score. **b. NOD EEG–fMRI correspondence.** Animate/inanimate decoding in 19 paired participants. Left, signed scores and ordinary least-squares fit. Right, rank correspondence. Pearson r = 0.844, Spearman ρ = 0.342, biweight r = 0.294, and leave-one-participant-out Pearson range = 0.623–0.882. Image sets are shared across instruments within each person, but differ between people; the pairing therefore includes stimulus-set effects. Matched-budget and image-difficulty analyses are reported in Methods and Supplementary Methods. **c. Cross-instrument associations.** Corrected Pearson and Spearman estimates for NOD and Wakeman–Henson. Lines show participant-bootstrap 95% confidence intervals for Pearson correlations. Datasets are not pooled. **d. Simultaneous COGITATE recordings.** Left, locked variance decomposition into participant-general, participant-by-instrument, participant-by-target and unresolved residual components. The model uses participant–modality–target battery summaries, not a reliability-adjusted run-level decomposition. The display rounds the general and instrument-specific shares to 43% and 11%. Right, odd-run EEG against even-run MEG balanced accuracy (n = 95; displayed r = 0.42, ρ = 0.35); the reverse split gives r = 0.30 and ρ = 0.31. More precise Pearson values appear in Results. **e. Own-map and own-operator advantages.** Participant-crossing tests in NOD and Wakeman–Henson. Points are bootstrap means and lines are 95% confidence intervals. Near-zero advantages indicate that common visual organization transfers more consistently than person-specific matching. **f. Category-blind BOLD responses.** Strict cross-fitted NOD event-related analysis. Curves summarize lower and upper decoding strata. Shading marks the prespecified event-lock window. The right-hand result gives the association with held-out balanced accuracy and held prediction; the displayed model value is R² = 0.664, P = 0.00002. This measures event-locked BOLD observability, not a direct measure of vascular reactivity.

The common component survived separation of runs. Odd-run EEG correlated with even-run MEG at r = 0.419; even-run EEG correlated with odd-run MEG at r = 0.297. The combined state-plus-substrate model explained only R² = 0.094 of the common factor and failed absolute prediction across sites. Much of the shared participant structure therefore remained unexplained by measured behavior, physiology and anatomy. The multilevel model used participant-by-modality-by-task summaries, not a separately identified run-level state decomposition (Methods).

External paired cohorts gave a less uniform picture (Fig. 4a–c). In NOD, EEG and fMRI signed margins were strongly correlated by Pearson’s measure (n = 19, r = 0.844; bootstrap 95% CI, 0.283–0.951; intact-pairing P = 0.00040). Rank and robust estimates were much smaller: Spearman ρ = 0.342, Kendall τ = 0.205 and biweight r = 0.294. Ten-percent winsorization gave r = 0.717. Deleting one participant at a time gave Pearson correlations from 0.623 to 0.882; maximum Cook’s distance was 3.67. Approximate correction for reliability gave r = 0.938. The large Pearson association reflected the spacing of scores more than a common ordering throughout the sample. NOD EEG–MEG correspondence was moderate (r = 0.362; ρ = 0.375), and MEG–fMRI correspondence was weak (r = 0.119; ρ = 0.218).

NOD also linked each participant to a particular stimulus set. Its 30 fMRI participants viewed different sets of 1,000 images, reused for that person across modalities. The paired association therefore included both person and stimulus-set effects. Matching observation budgets retained EEG–fMRI margin covariance (n = 19, r = 0.771; ρ = 0.412). After adjustment for observation counts and an exploratory image-difficulty proxy trained in other participants, the matched-budget association fell to r = 0.289 (95% CI, −0.104 to 0.648; conditional permutation P = 0.235). For full-data balanced accuracy, the adjusted association was r = 0.042. The proxy cannot separate the causal contribution of stimuli, but the original r = 0.844 cannot be assigned solely to the person.

Wakeman–Henson showed a different pattern (n = 16). EEG–GRAD correspondence was positive (r = 0.434; ρ = 0.568), EEG–MAG was negative to null (r = −0.187; ρ = −0.056), and EEG–fMRI was null (r = 0.127; ρ = −0.053). Pearson correlations were 0.504 for GRAD–MAG, 0.556 for GRAD–fMRI and 0.598 for MAG–fMRI, but bootstrap intervals included zero and rank estimates were weaker. This dataset distinguishes intact faces, both famous and unfamiliar, from scrambled images. Equal-budget EEG–fMRI margins correlated at r = 0.362, whereas restricting the analysis to the same 430 images gave r = 0.020. Repeated filenames were grouped across runs; semantic aliases could not be fully checked without the original image bitmaps. These corrected results replace the earlier strong MAG–GRAD interpretation. A further candidate dataset, ds002814, gave a negative EEG–fMRI result (n = 21; r = −0.122; ρ = −0.121) and was retained as supporting rather than locked replication evidence.

One-factor measurement-error models also gave imprecise partitions (Extended Data). In NOD, the estimated shared fractions of reliable variance were 1.00 for EEG (bootstrap interval, 0.305–1.00), 0.067 for MEG (0.001–0.547) and 0.655 for fMRI (0.073–1.00); covariance RMSE was 0.080. Wakeman estimates ranged from 0.051 for EEG to 0.688 for GRAD, but all intervals were broad and reached or approached 1.0; covariance RMSE was 0.145. These descriptive fits neither separate person from stable stimulus-set effects in NOD nor identify precise individual reversals or a universal factor.

Spatial maps transferred more consistently than person-specific advantages (Fig. 4e). Cross-fitted fMRI category maps predicted held EEG topographies in NOD (mean spatial r = 0.299; 18 of 19 positive) and Wakeman (mean r = 0.274; 16 of 16 positive). Corrected Wakeman projections were also positive for MAG and GRAD. Using a participant’s own map conferred little or no advantage: NOD mean = −0.026 (95% CI, −0.064 to 0.012), and Wakeman mean = 0.021 (−0.003 to 0.046). Own-operator advantages were similarly small: NOD 0.008 (−0.012 to 0.031), and Wakeman −0.004 (−0.045 to 0.032). Common visual organization transferred across instruments; matching maps and operators to the individual did not explain participant rank.

NOD supplied a separate fMRI result that did not depend on the category contrast (Fig. 4f). Event-locked BOLD magnitude or reliability predicted fMRI decoding in each of three locked analyses. The strict cross-fitted hemodynamic response function (HRF) analysis gave r = 0.859 for partial event-lock R² and held R² = 0.664 for a four-feature model. A three-way rotation gave r = 0.718 and HRF-block R² = 0.245. A later component analysis gave actual-event r = 0.718 and leave-one-out R² = 0.401; peak-amplitude r = 0.659 and R² = 0.334; shape-only R² = 0.022; and precision-only R² = 0.103. These are different estimators, not repeated estimates of one model. They relate decoding to the strength and reliability of the event-locked BOLD response. Without separate physiology, they do not isolate vascular reactivity from neural responsivity, hemodynamic transformation, estimation quality and scanner noise.

Some participant differences were shared across instruments, but no single ordering described the paired data. The represented distinction, the measurement method and their coupling to the individual all mattered.

## Discussion

The main result is a distinction between repeatability and universality. A participant’s decoding performance was repeatable within an instrument, but did not define a single rank that carried unchanged across instruments and contrasts. The differences had several sources. Simultaneous EEG–MEG revealed a shared participant component. Intracranial recordings linked decoding to local category selectivity. Anatomy altered the signal received by the sensors. The instrument then determined how these differences appeared in the measured response.

### A stable difference need not be an intrinsic trait

Reliability shows that an ordering recurs. It does not establish that the ordering belongs to the person independently of the measurement. The high split reliabilities leave substantial participant variation to explain. Regional and contrast controls show that this variation is not generic classifier aptitude. The simultaneous EEG–MEG correlations across disjoint runs establish shared participant structure beyond same-run fluctuations, while participant-by-task and participant-by-modality effects establish its limits. The unit of comparison is the person measured for a particular distinction with a particular instrument.

### Attention changes trial quality more than participant rank

Attention and physiology were relevant, but did not supply the main explanation. Behavior, pupil, gaze, blinking, autonomic measures and artifacts varied between participants and affected individual observations [10,11,13,14,22]. EOG contained category information, and nuisance control reduced absolute electrophysiological accuracy. Yet the ordering of participants changed little. Favorable prestimulus states and lagged attention also failed to rescue weak participants selectively. These findings argue against a simple compliance account.

Contamination and degradation remain different questions. A signal may carry category labels, or it may impair the neural measurement without carrying those labels. Motion can do the latter in fMRI [16,18,23,24]. The historical COGITATE fMRI estimates require further reconciliation, and the available run tables do not identify the exact share of momentary state in all modalities. Neither limitation changes the electrophysiological result: nuisance control left most stable ordering intact.

### Local selectivity and measurement geometry both matter

Participants with sharper face–object tuning in anatomically comparable visual cortex were easier to decode. The difference was chiefly a smaller nonpreferred response relative to overall visual responsivity. It survived anatomical and amplitude controls and a denominator estimated from training data alone. It was present at 500 ms and when the stimuli were task irrelevant. The local effect therefore did not require prolonged viewing or an explicit demand to attend to the contrast.

The association is a description of local responses, not a cellular mechanism. iEEG response energy does not identify inhibition, interneuron classes or cortical layers. Clinical coverage was nonrandom, only 29 participants entered the core analysis, cross-site prediction was asymmetric, and small-sample P/N models were poorly calibrated. Letter/false-font selectivity did not predict face–object decoding. The evidence supports a local, contrast-specific correlate, rather than a uniformly superior cortex.

Anatomy supplied a second route to stable decoding differences. Identical modeled sources produced different sensor fields because their distances and orientations relative to the instruments differed. Planar gradiometers showed the steepest distance dependence. Source–sensor coupling was reflected in measured category separation, which substantially reduced the gain–decoding association when included in the model. The anatomy of the measurement can therefore matter even when the cortical source is held fixed.

Preferred and nonpreferred responses should not be expected to look identical in every modality [25,26]. iEEG samples local field potentials; EEG and MEG sample mixtures of electromagnetic fields; fMRI samples hemodynamically transformed responses. The observed differences are consistent with these transformations, although independent cohorts and preprocessing also differ. GRAD attribution changed with adjustment, and the withdrawn COGITATE fMRI component analysis supplies no evidence for a universal component signature.

MEG made the distinction between total amplitude and useful information clear. Whole-head RMS did not predict participant rank. Category separation and variance along the discriminating axis did. Recurrence was strong within a contrast but did not generalize to an independent contrast. These measures describe the response geometry from which the decoder obtains information; because they summarize related distributions, their relation to decoding is not an independent upstream mechanism.

### Shared organization is not the same as shared participant rank

The external cohorts showed how a strong-looking cross-modal association can depend on its measure. NOD EEG–fMRI Pearson covariance was large and survived removal of any one participant, but rank and biweight correlations were modest. A small subset of score spacings contributed strongly without making the result a single-participant artifact. Moreover, each person carried the same private image set across modalities. Budget and difficulty controls reduced the association, so it cannot be identified as a person-only effect.

Wakeman gave a different pattern, with positive electrophysiology–fMRI Pearson correlations but broad intervals and weaker ranks. The candidate ds002814 result was null. These observations limit generalization without removing the shared structure seen in simultaneous COGITATE recordings. The one-factor models likewise allow shared and modality-specific variance, but their broad intervals and imperfect covariance fit do not permit a precise partition.

Common fMRI category maps predicted electrophysiological topographies, whereas a participant’s own map or operator conferred little advantage. Species-typical spatial organization can therefore transfer even when individual decoding rank does not. Larger paired samples, independent indicators within each modality, and participant-level errors-in-variables analyses will be needed to resolve that distinction fully.

### The practical difference is information per observation

Repeated observations narrowed the deficits of weak single-trial participants. Most did not lack the represented information; they supplied less recoverable evidence on each observation. GRAD category separation predicted accumulation efficiency in both directions of cross-site testing. Recurrence, variance, nuisance and forward gain added little once separation was known.

This changes the practical meaning of a weak recording. A participant who performs poorly under a fixed trial budget may be usable with more observations. An instrument that captures more information per trial can reduce recording time without changing the underlying representation. The distinction matters for experimental design, visual reconstruction and brain–computer interfaces [6,27,28,29]. The present curves do not specify reliable absolute trial requirements because their fitted asymptotes were unstable. Joint models across modalities and sessions, with better-constrained asymptotes, should address that design question.

### Consequences for multimodal studies

Disagreement between instruments need not be dismissed as noise around a single latent neural variable. It may be the result to explain. EEG, MAG, GRAD, iEEG and fMRI differ in spatial mixing, sensitivity and noise. Stable person-by-instrument effects are therefore expected. Mean rankings of modalities cannot determine which instrument will best measure a given participant and contrast, and reliability in one instrument cannot be transferred automatically to another.

Studies of individual decoding differences should report reliability, distinguish contamination from degradation and rank generation, fit predictors on separate data, and test absolute calibration as well as rank transfer. Negative held R² should remain visible. A person-general claim requires paired modalities and disjoint observations. A mechanistic claim requires a distinction between direct local recordings and reconstructed or hemodynamic measurements.

### Limitations

COGITATE EEG, MAG and GRAD were simultaneous, but iEEG and fMRI came from independent cohorts. External paired samples were small and differed in task and acquisition. Intracranial sampling was clinical; anatomy was incomplete for some participants; and no external intracranial cohort replicated the local result. Eye tracking and physiological channels were incomplete, and neither COGITATE nor external fMRI supplied audited separate physiology. The NOD event-lock measure therefore combines neural response magnitude, hemodynamic gain and estimation quality.

Cross-site calibration often failed even when rank transferred. Category separation and accuracy are mathematically close. The study is observational at the participant level and does not establish causal mediation or cellular mechanisms. External one-factor models did not support classification of individual discordance. Corrections to electrophysiological extraction and cache ordering, together with the unresolved historical all-trial fMRI measure, are documented in Methods. The clean fMRI benchmark reproduced exactly; the all-trial P/N branch was withdrawn, and inherited nuisance and upstream results remain unrevalidated. Candidate ds002814 supplies supporting negative evidence; ds003688 supplies no neural inference.

A participant is not simply easy or difficult to decode. The difference is stable, but it is a difference in what a particular representation and instrument make observable. Its practical expression is the amount of information recovered from each observation.

## Methods

### Datasets and participants

#### COGITATE M-EEG

The simultaneous M-EEG cohort comprised 100 participants from sites CA and CB in the COGITATE experiment [30]. Retained face–object decoding measures were available for 95 EEG participants and all 100 MAG and GRAD participants. Participants viewed faces, objects, letters and false fonts over five runs, with position, duration and task relevance varied experimentally.

Recordings included EEG, magnetometers, planar gradiometers, EOG, ECG-derived measures, valid pupil and gaze channels where available, behavior and artifact summaries. Individual anatomy and forward-model derivatives were available for most participants. Head translation was available for a subset.

#### COGITATE iEEG

Thirty-eight participants contributed intracranial recordings. Thirty-three entered the visual-contact decoding benchmark, and 29 participants with 569 visual contacts entered the core local-tuning analysis. Recordings combined electrocorticography (ECoG) and stereoelectroencephalography (SEEG) from clinical sites CE, CF and CG. Electrode coverage was determined clinically. Visual contacts were selected by anatomy, independently of decoding. Missing native anatomy and eye measures were not imputed.

#### COGITATE fMRI

The acquired cohort contained 118 participants; 102 entered the retained biological analysis at sites CC and CD. Analyses used visual-cortex trial patterns and image-derived nuisance measures. No separate physiological recording was treated as available. The historical N = 44 mechanism table and malformed signed margins were excluded. The clean common benchmark and historical all-trial measure are distinguished below.

#### External paired cohorts

NOD supplied paired EEG, MEG and fMRI recordings [31]. Modality overlap gave n = 19 for EEG–MEG, n = 19 for EEG–fMRI and n = 30 for MEG–fMRI. MEG participant sub-07 was reconstructed from a repaired, non-zero cache. Wakeman–Henson supplied EEG, MAG, GRAD and fMRI for 16 participants after correction of electrophysiological extraction [32]. Candidate ds002814 supplied paired EEG–fMRI for 21 participants and was used only as supporting negative evidence. Dataset ds003688 was not used for neural inference because only its anatomy was verified in the package.

### Input checks and corrections

We inspected raw and near-raw inputs visually and quantitatively. Checks covered finite values, variance, channel and time axes, baseline behavior, event-locked responses, channel positions, planar pairs, extreme amplitudes, cache provenance and sensitivity to individual participants.

M-EEG caches store all channels within each time bin and are therefore time-major. We reconstructed sensor-by-time arrays as:

`flat.reshape(n_trials, n_times, n_channels).transpose(0, 2, 1)`

Earlier direct reshaping to trial-by-sensor-by-time arrays was incorrect; all dependent values were replaced. For stimulus-locked electrophysiological displays, we used the RMS of the evoked sensor pattern rather than averaging signed sensors, which can cancel.

Wakeman–Henson and ds002814 electrophysiology were re-extracted with −0.5 to 0.6-s epochs, a 40-Hz low-pass filter and baseline correction. The defective short-epoch high-pass operation was omitted. NOD sub-07 MEG was rebuilt from valid input. Figure preparation used corrected channel order, actual montage metadata and pair-combined planar gradiometers.

### Primary decoding measures

The September replay reproduced all 430/430 common-benchmark participant measures from retained feature arrays. This was an array-level replay, not a new raw-preprocessing run. The following specifications define the analyzed measures.

#### EEG, MAG and GRAD extraction

We used five minimally preprocessed 250-Hz run derivatives per participant, extracted with `run_cogitate_comprehensive_decoding_battery.py`. Posterior channels were those with the smallest first-run head-coordinate y values: 24 EEG, 40 MAG and 80 GRAD channels. Ordered channel names were fixed across runs and recorded in the input manifests. MEG coordinates were transformed from device to head space. Selection used sensor coordinates rather than category responses. The EEG derivatives were average-referenced, bad channels were interpolated, and existing projectors were applied. The extractor retained the resulting channel lists rather than dropping annotated channels separately in each run.

Epochs extended from −0.2 to 1.6 s. We applied constant detrending, projectors, fourth-order Butterworth 1–40-Hz IIR filtering and a −0.2 to 0-s baseline correction. Decoder inputs were 25 non-overlapping 20-ms means from 0 to 0.5 s. Flattened arrays stored all selected channels in bin 1, then all channels in bin 2, and so on. The inverse was:

`flat.reshape(n_trials, 25, n_channels).transpose(0, 2, 1)`

Native amplitude units were preserved until training standardization.

The category-blind artifact gate used the largest channel peak-to-peak amplitude over the full epoch. We log10-transformed this value and rejected trials above the median plus 6 × max(1.4826 MAD, 1e−12). Trials had to pass every available sensor-family gate. This threshold used the participant’s whole session, rather than only classifier-training trials: it was label-blind, transductive quality control. The primary benchmark excluded target and response trials. Manifests specify the retained run, category, identity and cache-row mappings. Five participants lacked an eligible EEG family, giving n = 95 for EEG and n = 100 for MAG and GRAD.

#### Direct iEEG extraction

Inputs were `D:/cogitate/ieeg_all_category_binned_cache/sub-*_all_categories.npz`, produced by `decode_cogitate_ieeg_region_complementarity.py` and `decode_cogitate_ieeg_all_implants_roi.py`. ECoG and SEEG contacts required a known anatomical assignment that was not outside the brain. The visual mask matched lateraloccipital, cuneus, pericalcarine, lingual, fusiform, inferiortemporal or parahippocampal region names. It was independent of category decoding.

We retained the acquisition reference. The extractor imposed neither a new common-average reference nor a bipolar reference, so replay did not test robustness to alternative references. Target and hit/false-alarm trials were removed when caches were created. Complete epochs required data from −0.5 to 0.8 s. Fourth-order Butterworth 1–40-Hz second-order-section filters were applied forward and backward, followed by subtraction of the −0.2 to 0-s mean. Twenty-five means covered 0–0.5 s. Flattening was contact-major, unlike M-EEG.

Trials were rejected when median contact peak-to-peak amplitude in the analysis interval exceeded the median plus eight times the robust MAD scale. Native voltage units were retained. Numeric run/block labels were parsed from the first digit sequence. The anatomical benchmark contained 33 participants; the local-component analysis contained 29 participants and 569 contacts. These inclusion sets were kept separate.

#### fMRI common benchmark

For each participant, inputs were all_stimulus_trial_betas.npz (array lss_canonical) and all_stimulus_trials.csv in that participant’s directory under outputs/cogitate_fmri_lss_all_trials/. Source data comprised eight fMRIPrep native-space BOLD runs, confounds, FreeSurfer anatomy and event files. The visual region of interest used the extraction script’s anatomical labels on a 2-mm T1 grid. The eight run masks were intersected, with linear interpolation. Per-participant manifests retain voxel identifiers and image-grid provenance. No category-driven voxel mask was fitted outside training folds.

Voxel data were expressed as percent signal relative to the run mean. Voxels with near-zero means were zeroed using max(1e−6, 1% of the median absolute non-zero mean). Each onset–duration boxcar was convolved with:

h(t) = GammaPDF(t; 6) − GammaPDF(t; 16)/6.

The function was sampled at TR/10 over 32 s, normalized to unit sum, and then sampled at TR. Nuisance columns comprised 24 motion terms (parameters, derivatives, squares and squared derivatives), `a_comp_cor_00-05`, cosine terms, motion outliers and an intercept. Constant columns were dropped; the others were standardized. The same singular-value-decomposition nuisance projection was applied to BOLD and the trial design.

Least-squares-separate fits used one regressor for the target trial and the sum of the other trials. Designs were rejected when the smallest singular value was no greater than 0.01 of the largest; accepted fits used a pseudoinverse with `rcond=0.01`. Estimability was checked again after rejected trials were removed. Trial betas were centered within run and stored as float32. This category-blind, run-level estimation can leave shared residual structure among trials; held-out decoding does not make the trial estimates independent.

The clean common benchmark used face/object trials after exclusion of targets and hit/false-alarm trials. It reproduced exactly for all 102 participants. The historical all-face/object measure included target and response trials and estimated a different quantity. Matching that historical definition still left discrepancies in 24/102 participants, with maximum absolute difference 0.310997 balanced-accuracy units. Its component branch was withdrawn. All 102 participants had local preprocessed BOLD, confound, anatomy and event inputs, but the revision did not complete a fresh end-to-end BOLD extraction. This diagnosis replaces the earlier claim that repaired arrays were simply absent and the 0.269 discrepancy obtained by comparing different trial definitions.

### Feature selection and classifier

The primary feature cap was 64. Within training observations, we calculated the one-way ANOVA F statistic for each feature, set non-finite values to negative infinity, and selected min(64, number of features, number with finite F) using NumPy `argpartition`. Released fold manifests record selected indices, including ties. Each feature was centered by its training mean and divided by its training sample standard deviation (`ddof=1`); zero or non-finite standard deviations were replaced by one.

Let z be a standardized feature vector, μ_c the training mean for class c, and v the diagonal residual variance after subtraction of each training observation’s class mean (`ddof=2`). Variance was floored at 0.001 times the median positive variance. MAG and GRAD also used an absolute floor of 1e−12. The equal-prior diagonal discriminant score was:

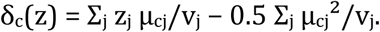

The predicted class had the largest score. Dividing by the positive training discriminant-vector norm defined the reported scores without changing the binary decision. MAG and GRAD used the equivalent midpoint binary score and a strict score > 0 rule. The common decoder sorted class names, face then object; the MAG/GRAD implementation coded face = 1. This distinction affects signed scores, not balanced accuracy. No off-diagonal covariance, tuned C, class weighting or estimated class prior entered this classifier.

### Separation of runs and identities

For each identity, we defined fold f as the integer represented by the first eight hexadecimal characters of SHA256(full identity string), modulo 5. For each run r and identity fold f:

test = (run = r AND fold = f); train = (run ≠ r AND fold ≠ f).

Both training classes and non-empty test data were required. MAG and GRAD additionally required at least 10 training observations per class and at least 10 same-run, other-identity observations for the archived calibration gate. The baseline classifier did not fit those calibration observations. Single-class test cells received predictions but no fold-level balanced accuracy. The common decoder required predictions for at least max(20, 70% of eligible trials).

Participant balanced accuracy pooled all estimable held-out predictions:

BA = 0.5[Σ_held face_ I(pred = face)/n_face_ + Σ_held object_ I(pred = object)/n_object_].

This was not an unweighted mean of fold accuracies. Neither a test run nor a test identity entered its corresponding classifier fit. Per-trial membership and per-fold feature selections permit exact reconstruction without regenerating random splits.

### Reliability and stable variance

We estimated participant-rank reliability from odd and even runs or partitions. Raw split correlations and Spearman–Brown corrected estimates were retained separately. Reliable between-participant standard deviation was the observed standard deviation multiplied by the square root of reliability. Face–object, four-category, position and duration measures tested content specificity without substituting a higher-capacity decoder. Regional controls compared posterior with anterior EEG, visual/core-posterior with nonvisual iEEG contacts, and visual/fusiform with frontal fMRI masks.

### Attention, behavior and physiological state

Category-blind features included task accuracy, median reaction time and its variability, omissions where available, relevance-condition behavior, pupil response and validity, gaze stability, blinks, EOG, heart rate and heart-rate variability, artifact rejection, and image- or movement-quality variables. Composite models were fitted within outer training-participant folds. Missingness handling, scaling, feature filtering, dimensionality reduction where used, and selection of regularization all used training participants only. Low-capacity ridge models were primary.

Participant-level models measured stable between-person associations. For valid run-resolved tables, we distinguished each person’s mean state from within-person deviations and same-run from lagged or cross-run prediction. The archived data did not contain a unified run matrix across modalities, so no omnibus within-person variance fraction was estimated. Retained analyses of prestimulus state, attention, relevance, duration and latency supplied the separate tests reported here.

### Contamination, degradation and participant ordering

We tested contamination by decoding category from auxiliary channels or nuisance-only features with runs and identities held out. We measured degradation by the change in neural balanced accuracy after nuisance regression fitted in training folds. We tested whether nuisance explained participant ordering by correlating raw and controlled scores and by predicting held-out participants from nuisance measures. Both linear and nonlinear controls were estimated in training data and applied to test data. fMRI motion-only decoding was treated separately from loss of neural decoding after motion and image-nuisance control.

### Nested prediction of participants

The upstream ridge model minimized:

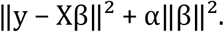

X contained an explicit constant, acquisition/site columns and retained family scores. The archived implementation used `fit_intercept=False`, so the constant and acquisition coefficients were penalized. Within every inner and outer training set, we retained features with at least 60% finite entries, median-imputed missing values, removed variance ≤ 1e−12, standardized features, and reduced each multifeature family to one principal-component score. Single-feature families were not reduced.

Outer folds were site-stratified: 10 folds, or 5 for n < 45, capped by the smallest site count; seed = 20260821. Inner site-stratified validation used four folds and seed = 20260821 + 100 + outer-fold index. We selected α by summed inner-validation mean squared error over 13 logarithmically spaced values from 1e−2 to 1e4. Every weight and transformation was learned within its training split.

For out-of-fold predictions, we calculated:

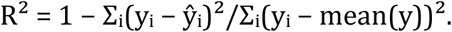

Model increments compared the same participants and folds. Calibration fitted y_i = a + bŷ_i. Pearson and rank correlations were reported separately from R². Cross-site models fitted all transformations and tuning on the training site alone.

The independent P/N reconstruction used a different specification: five shuffled outer KFold splits (seed 20260825), four inner folds (seed 20260826), nine logarithmically spaced α values from 1e−2 to 1e2, standardized P/N, site dummy columns and an unpenalized intercept. The corrected script, `recompute_strict_nested_models.py`, refitted scaling in every inner fold. It replaced an audit implementation that scaled the full outer-training set before inner validation. That earlier implementation did not expose outer-test observations, but inner validation was not fully isolated. Strict P + N held R² values were 0.115604 for EEG, 0.231618 for MAG, 0.175098 for GRAD and −0.030350 for iEEG. They are absolute prediction scores, not gains from recovery after a poor baseline.

### Variance decomposition

The archived simultaneous M-EEG model fitted participant-by-modality-by-target battery summaries, rather than repeated run-level indicators of the primary 64-feature benchmark. With chance_t denoting chance accuracy for target t, we defined:

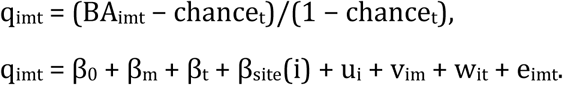

The random terms were independent, zero-mean Gaussians with variances σ_P², σ_PM², σ_PT² and σ_e². Statsmodels MixedLM used restricted maximum likelihood and L-BFGS, with `maxiter=500`. Each variance proportion was divided by their sum. The model did not separately identify run/state variance or a person-by-site effect and did not adjust for reliability. Its residual included unresolved higher-order interactions, state and error.

The battery used StandardScaler (`ddof=0`), training ANOVA selection of 128 features for native dimensions ≤ 800 and 256 otherwise, and `LogisticRegression(C=0.1, class_weight=’balanced’, solver=’lbfgs’, max_iter=1000, random_state=42)`. Cache identities were assigned by (numeric stimulus ID − 1) modulo 5. Historical split versions were not equated with the primary benchmark. The archived Methods record gives participant-general and participant-by-modality shares of 42.9% and 11.1%, respectively. These are model-specific battery variances, not fractions of reliable primary-benchmark variance. The ratio obtained after removing target and residual components is not the total reliable shared fraction.

External one-factor models used standardized odd/even signed-margin indicators, separately within each dataset:

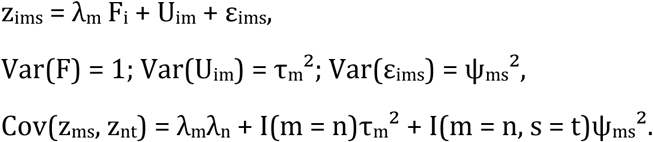

Positive variances were exponentiated parameters. Unweighted least squares fitted the upper covariance triangle with 12 starts, seed 20260818 and `max_nfev=4000`. We used 1,000 participant bootstraps with two starts. The shared reliable fraction was λ_m²/(λ_m² + τ_m²). These are descriptive covariance models. In NOD, the stable stimulus set is also linked to the participant.

### Direct iEEG local selectivity

Contacts were selected anatomically. Preference and face–object tuning were estimated in training runs and identities and evaluated on disjoint observations. We reversed the direction and averaged. Participant summaries included mean or top-quantile tuning, tuning normalized across all visual responses, preferred and nonpreferred responses, selectivity, recruited-contact summaries, and accumulation of contacts ranked in training data. Held-out observations never defined category preference.

Anatomical adjustment used Destrieux parcels, with complementary random-intercept and matched-contact resampling analyses. Available covariates included site, observation count, visual-contact count, ECoG/SEEG sampling, generic response reliability, category-signal variance and within-category variance. CE-to-CF and CF-to-CE predictions kept the test site separate. The amplitude sensitivity analysis removed contacts whose category-blind amplitude exceeded ten times the participant median before recomputing summaries. The same disjoint procedure was applied to task-relevance conditions and 500-, 1,000- and 1,500-ms durations. Nonpreferred recruitment refers to a response measure, not inhibition.

### Preferred, nonpreferred and selective responses

Training category-contrast norm selected the top ceil(20% of eligible elements); the category with the larger training response norm defined preference. One partition used odd runs with identity folds 0–2 and the other even runs with folds 3–4. Directions were reversed and averaged. iEEG used SHA256 folds. The locked M-EEG P/N calculation used numeric cache folds and retained task-irrelevant trials through its relevance parser. These partitions differed from the full run-by-hash decoder folds; the decoding measure did not come from an entirely independent recording.

EEG and MAG used sensors, GRAD concatenated both orthogonal planar components at each location, and iEEG used visual contacts.

For each selected element, p and n were the held-out preferred and nonpreferred category-mean time-course norms, and a was the category-blind all-visual evoked norm. P and N were the averages of p/a and n/a, with a floor of 0.05 times median a and numerical floor 1e−12, followed by averaging of directions. Sensitivity analyses kept the same selections and splits, but used no denominator, estimated a only from training data, or fitted the participant-level model:

zBA = intercept + β_P_ zP_absolute_ + β_N_ zN_absolute_ + β_A_ zA + site terms.

The joint model used either held-out or training A. HC3 intervals described regression uncertainty. Univariate intervals used 5,000 site-stratified participant bootstraps. New sensitivity P values were parametric and were kept separate from the original permutation P value.

For iEEG (n = 29), absolute N gave r = −0.136824 (P = 0.479), while training-normalized N gave r = −0.579001 (ρ = −0.684729; bootstrap 95% CI, −0.738666 to −0.462543; P = 0.000999). Absolute P gave r = 0.316825 (P = 0.094). Controlling training magnitude in the joint model gave absolute N β = −0.899585 (HC3 95% CI, −1.287771 to −0.511399) and P β = 0.00817. The association is therefore with nonpreferred expression relative to overall responsivity. Training normalization addresses reuse of the held-out denominator; it does not establish a cellular mechanism. Normalization CSVs retain all values, influence measures and directions.

### MEG response geometry

We estimated category-centroid separation, within-category variance parallel and orthogonal to the training discriminating axis, residual effective rank, temporal-profile recurrence, four-category subspace recurrence and noise-covariance recurrence from disjoint runs. Axes, covariance estimates, time profiles and normalizations were defined in training data. Global RMS was category blind. Temporal recurrence used the corrected sensor–time reconstruction. Noise-covariance recurrence was assessed without orienting its sign to the decoding measure.

Geometry models excluded exact decoder margins and were tested in both directions across sites. Attenuation analyses measured the change in reproducibility–decoding or gain–decoding associations after adding separation; they were not causal-mediation analyses.

### Forward models and head position

We calculated visual forward gain from each participant’s anatomy and lead fields using the fixed generic visual source model. Source–sensor distance, head geometry and sensor family were treated as properties of the measurement. Run-specific head-position models estimated within-session gain changes and predicted run-local category separation and held-run decoding.

Functional map/operator crossing used maps estimated in training data and held-out topographies. We compared each participant’s own map and operator with other or common maps and operators. These comparisons tested measurement organization; they were not neural perturbations.

### fMRI motion and category-blind event responses

COGITATE nuisance features included framewise displacement, DVARS, outlier burden and retained tissue/global summaries. Nuisance-only decoding, change in neural accuracy after control, and preservation of participant ordering were treated separately.

NOD event-lock analyses measured all-visual event-related BOLD responsivity without using animate–inanimate category labels. Three locked implementations were retained: a strict cross-fitted HRF model, a three-way rotation model, and a later summary-level component counterfactual. Their estimates were reported separately. Peak magnitude, temporal shape and precision were not measures of vascular physiology; no separate vascular or physiological measurements were available.

### Information accumulation

We fitted participant learning curves from repeated trial/identity aggregation with both runs and identities held out. The primary efficiency measure was negative log τ. Asymptotes and observations-to-criterion were retained in the audit record rather than used as fixed acquisition requirements. Cross-site models tested category separation, recurrence, parallel variance, nuisance and forward gain as predictors of accumulation efficiency. Displayed curves did not extrapolate participants beyond their available observations.

### External paired decoding and stimulus controls

NOD distinguished animate from inanimate images. Wakeman–Henson distinguished intact faces, famous and unfamiliar, from scrambled images. Neither used the COGITATE face–object contrast. Training standardization used sample standard deviation. Features were ranked by squared class-mean difference divided by the mean class-specific standardized sample variances, with floor 1e−8; the top 64 were retained. The diagonal discriminant used the class midpoint and divided scores by the training weight norm. Both training classes required at least two observations; single-class test cells were skipped. Runs and SHA256 identities were jointly excluded. This classifier differs from the common ANOVA/residual-variance classifier.

The observation inventory lists counts by participant, modality, run and class. NOD typically contained 4,000 EEG/MEG observations and 1,000 fMRI images, with 32 EEG, 20 MEG and 10 fMRI runs. The 30 fMRI participants viewed disjoint sets of 1,000 images, giving a union of 30,000 images. Each person’s images recurred across modalities. Intact-pairing permutations therefore tested correspondence of the person plus the linked image set, not the person alone. Reliability correction does not remove this confound.

Matched-budget analyses selected the first 10 NOD runs and 26 trials per class per run, for 520 trials, or the first six Wakeman runs and 28 trials per class per run, for 336 trials. Deterministic SHA256 selection used ten seeds. We reran decoding after selection without changing training-only transformations or run/identity exclusion. Participant scores were averaged over seeds. Intervals resampled participant pairs, not seeds as independent participants. The trial-selection manifest is released.

Wakeman repeated filenames were grouped across runs before splitting. EEG, MAG and GRAD shared 448 filename identities across participants; fMRI shared 432. Their common intersection contained 430 identities. One sensitivity analysis retained one presentation of each common identity per modality. Additional split sensitivities standardized case and leading zeros and conservatively grouped numeric families across famous, unfamiliar and scrambled prefixes. These procedures do not prove semantic equivalence. Original stimulus bitmaps were not locally available for content-hash or alias checks.

NOD fMRI’s non-overlapping stimulus sets prevented direct leave-one-person-out estimates of difficulty for the same image. We therefore fitted an exploratory proxy using fixed low-level and frozen ResNet18 features to predict correctness in other participants. Ridge regression used α = 100 and training standardization. Every target-participant image was excluded from fitting. Mean predicted ease over the target participant’s full image set defined the difficulty proxy.

Partial correlations adjusted modality scores for observation counts and the two modality-specific proxies. The proxies were averaged over full image sets, not over each selected matched-budget subset. Bootstraps and 10,000 residual-pairing permutations conditioned on the estimated proxies rather than refitting the full feature-to-difficulty pipeline. These were exploratory sensitivity analyses, not a confirmatory person-factor estimate.

For NOD EEG–fMRI (n = 19), full-data margins gave r = 0.843580 and ρ = 0.342105; equal-budget margins gave r = 0.770882 and ρ = 0.412281. After adjustment for counts and difficulty proxies, full-data margin r was 0.513205 and equal-budget margin r was 0.289073 (95% CI, −0.103979 to 0.647643; conditional P = 0.235176). Full-data balanced accuracy fell to partial r = 0.042373. For Wakeman (n = 16), equal-budget margin r was 0.361930 and identical-430-image margin r was 0.020391. Full estimates and seed sensitivities were retained separately by dataset.

### fMRI map projection and operator crossing

Category maps were fitted in training fMRI partitions and projected through electrophysiological sensor models. Predicted and observed held-out topographies were compared after unit-norm scaling when spatial correlation was the outcome. We checked EEG montage order, invalid channels and duplicate positions. Planar gradiometers were combined at unique pair locations, rather than interpolating co-located signed components. Group prediction was distinguished from advantages of a participant’s own map or operator. Cortical and scalp displays used validated metadata and common scales for direct comparisons.

### Statistical inference

Archived upstream intervals used 5,000 bootstraps of observed/predicted held-out participant pairs; they did not refit every model. The 10,000 pairing permutations conditioned on fixed predictions. Separately archived 100-permutation analyses refitted the full pipeline. These procedures test different null questions; fixed-prediction permutations are not fully nested refits. Negative held R² values were retained.

### Analysis provenance and exclusions

The analyses derive from the corrected evidence package dated 23 August 2026, subsequent structural analyses, and an independent reconstruction dated 25 August 2026. The September revision brought these records together without equating different model specifications. The analysis record distinguishes locked primary, locked supporting, corrected external, independently reconstructed, candidate/exploratory, superseded and unavailable results.

Outputs with incorrect feature order, the historical fMRI N = 44 mechanism analysis, malformed fMRI margins, the invalid COGITATE HRF pilot, unrepaired external caches and ds003688 neural claims were excluded from primary inference. Historical comparison values remain labeled in audit files. Corrections to external electrophysiology and M-EEG ordering were found during final quality control; reported external and P/N/selectivity results use corrected outputs.

The earlier 0.269-BA fMRI discrepancy compared clean and all-trial definitions. Matching definitions instead found discrepancies in 24/102 all-trial measures, with a maximum of 0.311, while every clean common-benchmark measure reproduced to numerical precision. The all-trial P/N branch was withdrawn. Historical fMRI nuisance and upstream estimates that inherited it remain unrevalidated supporting results. Local source inputs exist; missing raw data are not the stated cause. Valid NOD event-lock analyses are separate from these exclusions.

The revision supplies executable replay, extraction snapshots, model scripts, configurations and split manifests. It does not claim that every historical preprocessing step was independently rerun. Full raw-to-result regeneration and semantic-alias checking still require controlled source inputs and the original Wakeman stimulus bitmaps. A complete run-resolved trait–state decomposition and confirmatory participant-level Bayesian classification of discordance remain unavailable.

### Figure preparation

Original figures were prepared at 178-mm width with vector text and axes; only dense anatomy and large point clouds were rasterized. Scalp maps used the correct channel order and montage metadata, common scales within comparisons, and explicit color bars. Checks covered overlap, clipping, legend collisions, invalid interpolation, duplicated planar locations, grayscale legibility and simulated protanopia/deuteranopia.

Figure 1a uses an uninterrupted 26-s segment from the original 1,000-Hz same-run M-EEG recording, including EEG, MAG, planar GRAD, EOG, ECG, analog gaze and analog pupil. SOURCE_DATA_INDEX.csv and the project manifest map panels to their sources.

## Data and code availability

The accompanying paper package contains the manuscript, four figures, source-data index, citation audit and editorial change log. Analysis code, trial/fold manifests, source-data tables and numerical audits are supplied in the companion Methods and robustness package. Public external datasets retain their original access terms; dataset identifiers and access information are listed in the complete Methods and reference ledger. Release of COGITATE participant-level derivatives is subject to consortium approval and the applicable data-sharing agreements. Historical and failed estimates remain in explicitly labeled audit and comparison files, rather than serving as primary evidence.

